# Translation regulation of ATF4 by the termination complex Hbs1-Pelo is required for visual system development and function

**DOI:** 10.1101/2025.10.08.681214

**Authors:** Katherine Querry, Inês Lago-Baldaia, Narayanan Nampoothiri V.P., Christopher Garbark, Abby J. Carney, Vilaiwan M. Fernandes, Deepika Vasudevan

## Abstract

Inherited retinal diseases are a class of genetically heterogenous disorders characterized by mutations in genes required for retinal function, resulting in progressive loss of vision in human patients. One such deletion mutation in the translation termination factor, HBS1L, results in a suite of developmental anomalies in human patients, including progressive vision loss defects. HBS1L, and its interaction partner Pelota (Pelo), are required for recycling stalled ribosomes on mRNAs, but the specific mRNA target causative of vision defects seen with HBS1L deletion remains unknown. Further, the specific cell types in the visual system that require HBS1L for proper development and function are also unknown. Here, we discover that loss of the *Drosophila* HBS1L homolog, *Hbs1*, results in reduced expression of the stress responsive Activating Transcription Factor 4 (ATF4) which is encoded by an mRNA containing multiple upstream open reading frames (uORFs) in its 5’ leader. We corroborate these results in cultured human cells, where we find that HBS1L and Pelo promote translation reinitiation at the ATF4 ORF by facilitating proper translation termination at the preceding uORFs. Like human HBS1L deficiency patients, loss of function *Drosophila* mutants for *Hbs1*, *pelo*, and *ATF4* show vision defects as measured by electroretinograms (ERG). Depleting *Hbs1* in lamina neurons replicated the ERG defects seen in *Hbs1* mutants, suggesting that Hbs1-Pelo is required for proper lamina neuron function. Further confocal analysis of *Hbs1* mutants revealed ‘vacuolization’ defects in the lamina layer, which are indicative of defective synapse formation between lamina neurons and photoreceptors. Strikingly, restoring ATF4 expression in the lamina partially rescued ERG defects in *Hbs1* mutants, indicating that ATF4 is likely a relevant mRNA target regulated by Hbs1-Pelo in these cells. Together, our data support a model wherein Hbs1-Pelo mediated translation regulation of ATF4 in lamina neurons underlies the inherited retinal disease caused by HBS1L deletion.

## Introduction

Organisms have evolved an array of stress response pathways to counteract a wide spectrum of extrinsic and intrinsic stressors. In higher organisms, these protective mechanisms have been co-opted to play essential roles in normal tissue development and homeostasis, ensuring that cells adapt and function optimally under varying physiological conditions. A classic example of such a mechanism is the Integrated Stress Response (ISR), an evolutionarily conserved pathway initiated by stress responsive kinases. ISR signaling famously regulates the translation landscape by reducing global translation by limiting initiator methionine availability (Ryoo and Vasudevan 2017). These limiting conditions however promote the translation of stress responsive mRNAs with a specific 5’ leader architecture, ATF4 (Activating Transcription Factor 4) being a prime example (Ryoo and Vasudevan 2017). Loss of ATF4 has been linked to a plethora of developmental abnormalities in the skeletal system, hematopoiesis, adipose and liver tissues, and the visual system. While we have a fair understanding of many ATF4 phenotypes, the role of ATF4 in visual system development is relatively understudied.

The lens in the eye focuses light onto the retina, which is comprised of several types of highly specialized neurons that are organized in distrinct stratified layers. Phototransduction is initiated in photoreceptors where light is sensed by opsin proteins which trigger a change in membrane potential. This change in membrane potential modulates neurotransmitter release, sending visual information to downstream neurons. Photoreceptors form synapses with bipolar neurons (Sanes and Zipursky 2010), amongst others, which then transduce the signal on to ganglion cells, whose axons form the optic nerve. Several studies have shown that loss of ATF4 results in developmental abnormalities in the lens in mice (Nandakumar et al. 2025). ATF4 expression is also robustly detected in developing photoreceptors (Ooe et al. 2017; Kang et al. 2015; Vasudevan et al. 2022; Preston et al. 2025) and is also elevated in the aging retina (Ooe et al. 2017; Vasudevan et al. 2022). In addition to its role in lens development in mice, we previously demonstrated that *ATF4* mutants display age-dependent retinal degeneration in *Drosophila* (Vasudevan et al. 2022). ATF4 has also been implicated in the progression of many retinal degeneration diseases including autosomal dominant retinitis pigmentosa, diabetic retinopathy, and Fuchs’ endothelial corneal dystrophy amongst others (Bhootada et al. 2016; Pitale et al. 2017; Vasudevan et al. 2022; Qureshi et al. 2024). Given these profound developmental and clinical implications, there is significant interest in identifying novel regulators of ATF4 and delineating the specific visual system cell types critically dependent on ISR signaling for their normal development and function.

The ATF4 mRNA is almost ubiquitously detected in mice and *Drosophila* tissues (Leader et al. 2018, 2; Yang and Karsenty 2004), with ATF4 protein levels being regulated primarily at the level of mRNA translation (Neill and Masson 2023). Across phyla, the 5’ leader sequence of ATF4 mRNA contains upstream open reading frames (uORFs) in addition to the main open reading frame which encodes ATF4 protein (Hinnebusch et al. 2016). While the number of these uORFs varies across species, the general regulatory architecture is evolutionarily conserved from yeast to *Drosophila* and humans (Hinnebusch 1984; Lu et al. 2004; Vattem and Wek 2004; Kang et al. 2015). Human ATF4, for instance, possesses two uORFs, with the second uORF partially overlapping the ATF4 coding sequence itself (Lu et al. 2004; Vattem and Wek 2004). It is worth noting here that there are discrepancies in the number of annotated uORFs in the human ATF4 5’ leader, with some studies regarding there to be only two uORFs (Vattem and Wek 2004; Lu et al. 2004), since they exclude a zero-length (start-stop) ORF that is most distal to the ATF4 start codon (Bohlen et al. 2020; Rendleman et al. 2024; Smirnova et al. 2024). Under homeostatic conditions, translation typically initiates and terminates at uORF1, with only a subset of ribosomes reinitiating at uORF2 (Hinnebusch et al. 2016). However, due to the overlapping nature of uORF2 with the ATF4 coding region, ATF4 protein synthesis does not occur when initiator methionyl-tRNA bound to the initiation factor eIF2 is abundant. Upon stress-induced phosphorylation of the alpha subunit of eIF2 by ISR kinases, the resulting reduction in active eIF2 complex abundance leads to ribosomes skipping initiation at uORF2. Consequently, ribosomes instead reinitiate translation at the main ATF4 ORF, resulting in increased ATF4 synthesis during ISR activatoin. In addition to being regulated by eIF2 availability as modulated by ISR kinases, translation initiation at the ATF4 ORF has been shown to be dependent on specific initiation factors (Roy et al. 2010; Herrmannová et al. 2024) and 5’ leader features (Chan et al. 2013; Rendleman et al. 2024; Smirnova et al. 2024). However, efficient termination at uORF1 is also a key regulatory step in reinitiation at the ATF4 ORF (Ait Ghezala et al. 2012; Bohlen et al. 2020; Vasudevan et al. 2020).

Termination commences with stop codon recognition, followed by release factor recruitment, peptidyl-tRNA hydrolysis, and finally ribosome dissociation. High resolution ribosome profiling data have revealed that the tRNA binding factors, DENR-MCTS1 heterodimer and its homolog eIF2D, efficiently remove spent tRNAs from the penultimate codons in uORF1 (Bohlen et al. 2020). Such removal of spent tRNAs is predictably crucial for proper termination at uORF1, and loss of these factors significantly impaired reinitiation at the ATF4 ORF (Bohlen et al. 2020). These results raise the intriguing possibility that there may be additional such termination factors which regulate other critical steps of translation termination at uORF1. Our work herein substantiates this possibility by discovering that the ribosome recycling factor, HBS1L and its binding partner Pelo, are required for efficient ATF4 translation. We use a *Drosophila* model to describe that such regulation is crucial for proper functioning of the visual system. Importantly, our work proffers ATF4 as a mechanistic target underlying the vision defects and other developmental anomalies seen in human patients with HBS1L deficiency.

## Results

### Identifying Hbs1-Pelo as regulators of Drosophila ATF4

Recent work demonstrated that loss of the tRNA-binding protein complex, DENR-MCTS1, results in the accumulation of 40S intermediates at certain penultimate codons, including in the uORF1 of ATF4 mRNA (Bohlen et al. 2020). Based on this, we hypothesized that the action of ribosome recycling factors, whose function is to separate the 40S and 60S subunits, precedes removal of spent tRNA DENR-MCTS1/eIF2D. Since we had previously used the *Drosophila* model to discover that loss of DENR-MCTS1/eIF2D resulted in reduced ATF4 translation, we performed a targeted RNAi screen to test whether known termination and ribosome recycling factors were required specifically for ATF4 translation.

In eukaryotes, the release factor eRF1 (eukaryotic release factor 1) binds to the stop codon, and eRF3, a GTPase, facilitates the release of the newly synthesized polypeptide from the ribosome (Hellen 2018). The eRF1 and eRF3 proteins subsequently dissociate from the ribosome, and the ribosomal subunits (40S and 60S) are separated by the ATP-binding cassette protein ABCE1 (Hellen 2018). In addition to these core termination factors, a host of other factors with paralogous function have been identified. Pelota (Pelo) has been shown to substitute for eRF1 function (Atkinson et al. 2008; Pisareva et al. 2011; Hellen 2018). In mammals, two distinct genes encode two different eRF3 forms, namely eRF3a and eRF3b (Chauvin et al. 2005). The GTPases HBS1L (Hsp70 Subfamily B Suppressor 1-like), GTPBP1 and 2 (GTP-binding protein 1, 2) have likewise been shown to perform functions similar to eRF3 (Atkinson et al. 2008; Pisareva et al. 2011; Ishimura et al. 2014; Terrey et al. 2020).

To study the effects of the above described release factors on ATF4, we employed a faithful reporter of ATF4 transcriptional activity, 4E-BP^intron^-DsRed, which is constitutively expressed in the wandering third instar larval fat tissue (also known as fat body) (Kang et al. 2016). We identified the *Drosophila* homologs of all release factor GTPases (Marygold et al. 2016) (**Table S1**) and utilized the GAL4-UAS system (Brand and Perrimon 1993) to RNAi deplete them in the fat body. Using a fat body driver, *Dcg-GAL4* (Asha et al. 2003) we found that two independent *UAS-RNAi* lines targeting *Hbs1*, the *Drosophila* homolog of HBS1L, led to reduced ATF4 activity in adipocytes in comparison to adipocytes expressing a control *UAS-LacZ^RNAi^* (**Fig. 1A, B**). We did not recover any *Dcg>eRF3^RNAi^*animals, and we also did not record appreciable changes in DsRed levels with the other RNAi lines tested. Notably, depleting *Hbs1* in the fat body did not influence the expression of a *UAS-GFP* transgene (**Fig. 1A**), indicating that the effects of Hbs1 were specific to ATF4. We validated these results in *Hbs1* loss of function mutant animals (transheterozygous *Hbs1^1^/Hbs1^48^* (Li et al. 2019)), which also showed a marked decrease in 4E-BP^intron^-DsRed in the fat body in comparison to wildtype *w^1118^* animals (**Fig. 1C, D, S1A**).

**Fig. 1.**
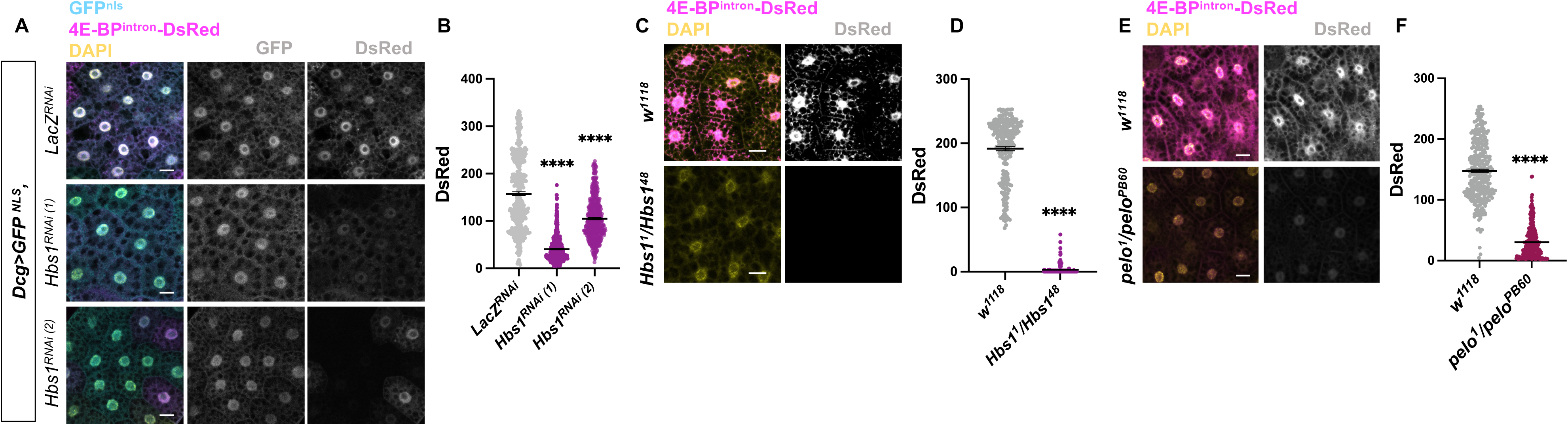
*Hbs1* and *Pelo* are required for ATF4 activity in the *Drosophila* fat body. **A**. Confocal images showing fat bodies from animals expressing an ATF4 reporter (4E-BP^intron^-DsRed, magenta) and GFP (cyan) driven by *Dcg-GAL4*, which also drives expression of the indicated *RNAi*. **B**. Quantification of DsRed intensity from A. Each data point represents one nucleus, and black bar represents mean of data from at least 5-7 animals across two independent crosses. Error bars represent standard error. Statistical comparisons were made using Kruskal-Wallis test with Dunn’s correction. **C, E.** Confocal images of fat bodies expressing 4E-BP^intron^-DsRed (magenta) from wildtype (*w^1118^*), and transheterozygous mutants for *Hbs1* (*Hbs1^1^/Hbs1^48^*), and *pelo* (*pelo^1^/pelo^PB60^*). **D, F.** Quantification of DsRed intensity from C and E respectively. Each data point represents one nucleus, and black bar represents mean of data from at least 5-7 animals across two independent crosses. Error bars represent standard error of mean. P-values were calculated using Mann-Whitney test. Scale bars in A, C, E: 25μm; Nuclei are counterstained by DAPI (yellow). ****=p<0.0001, ***=p<0.001, **=p<0.01, *=p<0.05 with only significant comparisons shown here and in future figures.

Next, we sought to identify whether Pelo, the release factor known to partner with Hbs1, is required for ATF4 activity in the fat body. Fat bodies from *pelo* loss of function mutant animals (transheterozygous *pelo^1^/pelo^PB60^* (Yang et al. 2015) showed a marked decrease in 4E-BP^intron^-DsRed (**Fig. 1E, F, S1B**), similar to what we observed with loss of *Hbs1*. We also sought to test the other known release factor, eRF1. However, *Drosophila eRF1* loss of function mutants have been reported to be lethal (Chao et al. 2003) and we were also unable to recover animals when we knocked down *eRF1* in the fat body. Based on these results, we concluded that the Hbs1 acts together with Pelo to regulate ATF4 in *Drosophila*, and we proceeded to investigate whether such regulation was conserved in vertebrates.

### HBS1L-Pelo regulate ATF4 translation by promoting efficient termination at the upstream ORF

To test whether HBS1L-Pelo regulate ATF4 in human cells, we siRNA-depleted these factors in HEK293T cells. To robustly detect ATF4, we treated cells with the ER stress-inducing chemical, Tunicamycin. Western blot analyses revealed that the siRNA-depletion indeed led to reduced HBS1L and Pelo protein in cells (**Fig. 2A**). While cells transfected with control siRNA showed nearly five-fold induction in ATF4 with ER stress, depleting either HBS1L or Pelo resulted in significantly lower ATF4 levels (**Fig. 2A, B**). These results strongly suggested that HBS1L-and Pelo-mediated ATF4 regulation is conserved between *Drosophila* and humans. Further, we observed no change in ATF4 mRNA levels in HEK293T cells with siRNA treatment (**Fig. S2A**), leading us to posit that HBS1L-Pelo likely regulate ATF4 translation.

**Fig. 2.**
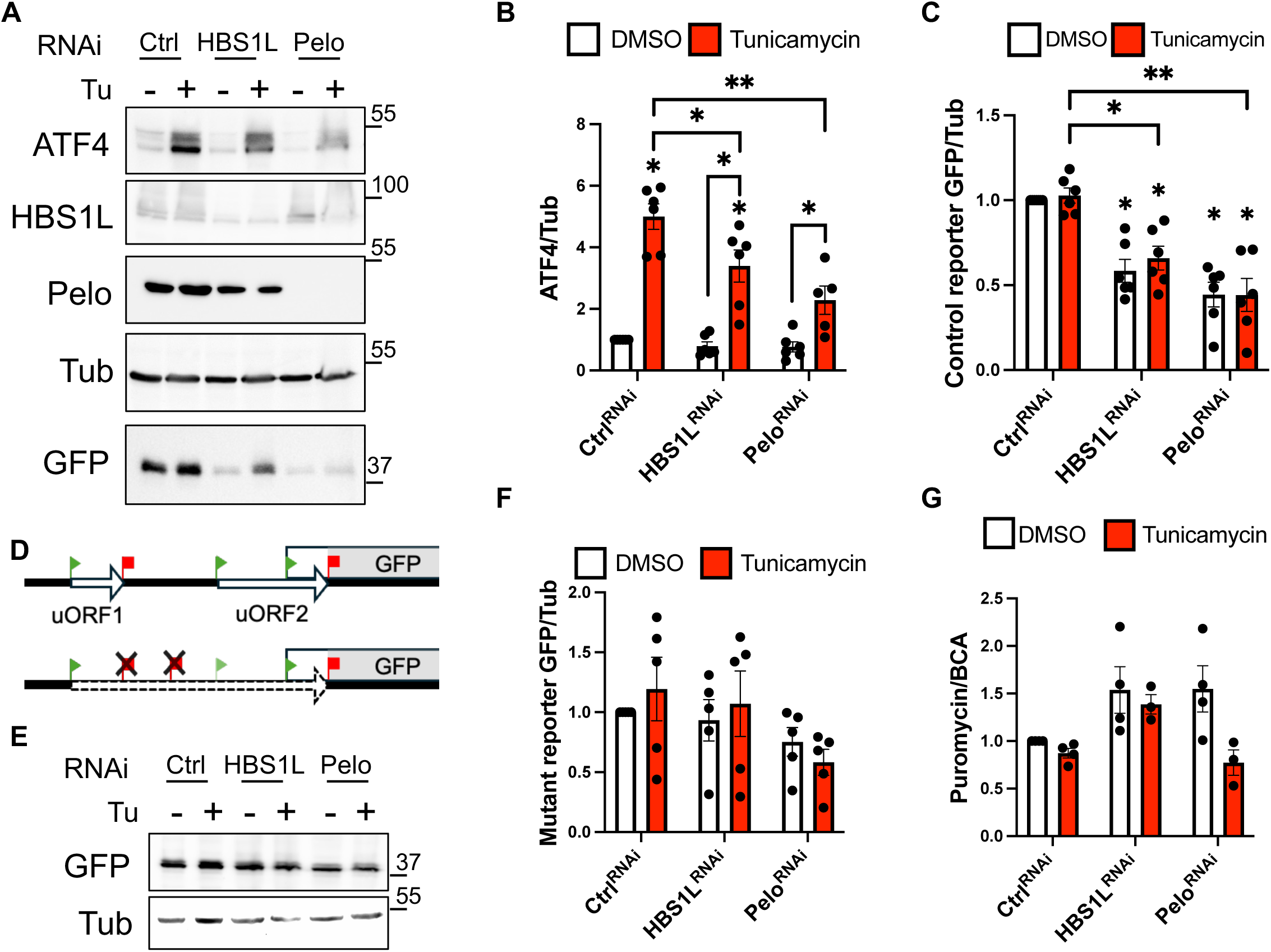
HBS1L and Pelo regulate ATF4 ORF translation by promoting efficient termination. **A.** Representative western blot analyses of lysates prepared from HEK293T cells co-transfected with siRNA (Ctrl (negative control), HBS1L, Pelo) and an ATF4 5’ leader-GFP reporter where the coding sequence of ATF4 is replaced with GFP (see schematic in D). Cells were treated with 10μg/ml Tunicamycin (Tu) for 4 h to induce ER stress prior to lysis. Tubulin (Tub) serves as a loading control. **B-C.** Quantification of ATF4 intensity (B) and GFP intensity (C) from (A) normalized to loading control (Tub). Data represent mean of at least five independent experiments; error bars are standard error of mean. **D.** (Top) schematic of control ATF4 5’ leader-GFP reporter, and (bottom) a mutant reporter where all the stop codons upstream of the ATF4 start codon are deleted. Green triangles mark start codons, red squares mark stop codons. Arrows in solid black outline represent open reading frames corresponding to uORFs. Dotted black arrow (bottom) represents the new putative reading frame generated by mutating stop codons in the 5’ leader. **E.** Western blot analysis of mutant ATF4 5’ leader-GFP in HEK293T cells where HBS1L and Pelo are RNAi depleted. **F.** Quantification of GFP signal from E normalized to Tub. Data represent mean of at least five independent experiments; error bars are standard error of mean. **G.** Quantification of Puromycin signal normalized to total protein (as measured by BCA) in HEK293T cells where HBS1L and Pelo are RNAi depleted. Data represent mean of four independent experiments; error bars are standard error of mean. See S2B for representative blot. Statistical comparisons with Ctrl^RNAi^-DMSO were performed using the Wilcoxon signed rank test for and are shown as floating asterisks when significant. Comparisons with Ctrl^RNAi^-Tunicamycin were performed using the Friedman test with Dunn’s correction. Other comparisons were performed using the Wilcoxon matched pairs signed rank test.

ATF4 translation is heavily regulated via the 5’ leader of the ATF4 mRNA. To test whether HBS1L and Pelo act on the 5’ leader of ATF4 mRNA, we utilized a reporter where the ATF4 5’ leader is placed upstream of GFP (Lu et al. 2004) (see schematic in **Fig. 2D**). Consistent with our hypothesis, we observed that knockdown of either HBS1L or Pelo resulted in reduced ATF4 5’-GFP reporter expression under both vehicle-treated and ER stress conditions (**Fig. 2A, C**). We’d like to note here that consistent with previous reports, the ATF4 5’-GFP reporter does not show appreciable inducibility with ER stress unlike endogenous ATF4 even in control cells. We attribute to this to the perdurance of GFP which is proteostatically more stable in comparison to endogenous ATF4 (Lu et al. 2004). Nonetheless, our results demonstrate that HBS1L-Pelo regulate translation of ATF4 via the 5’ leader region.

Since the best studied role for HBS1L-Pelo is in ribosome recycling (Atkinson et al. 2008; Pisareva et al. 2011), we further hypothesized that HBS1L-Pelo likely facilitate reinitiation at the ATF4 main ORF by promoting efficient termination at stop codons preceding the ATF4 start codon. Such efficient termination would allow for DENR-MCTS1/eIF2D to effectively remove spent tRNAs and generate reinitiation-competent 40S ribosomes. If this model is correct, eliminating all termination events preceding reinitiation at the ATF4 start codon should eliminate the effects of HBS1L-Pelo on ATF4 translation. To test this, we used a mutant version of the ATF4 5’-GFP reporter wherein all stop codons in the 5’ leader are mutated (Lu et al. 2004) (**Fig. 2D**). Consistent with our model, we observed that siRNA knockdown of HBS1L did not impact expression of the mutant reporter (**Fig. 2E-F**). Interestingly, knockdown of Pelo resulted in a small but statistically significant reduction in the mutant reporter. This could be because loss of Pelo results in a global decrease in translation, which we tested using a puromycin incorporation assay to measure global translation rates in HBS1L and Pelo knockdown cells. Our data showed that depleting either HBS1L or Pelo did not result in a statistically significant change in overall translation rates under vehicle-treated or ER stress conditions (**Fig. 2G, S2B**). However, we did observe that the relative decrease in translation rates between vehicle-treated and ER stress conditions was much more pronounced in Pelo siRNA cells in comparison to control siRNA cells (**Fig. 2G, S2B**). This analysis suggests that the decrease in ATF4 protein seen with Pelo depletion in **Fig. 2A-B** may be partly due to a decrease in global translation seen with loss of Pelo. Nonetheless, our combined results support a model wherein HBS1L-Pelo mediate efficient translation termination at uORF1 and thus promote reinitiation at the ATF4 ORF.

### Hbs1 mutants exhibit phototransduction defects

Our data thus far showed Hbs1 and Pelo to be regulators of ATF4 in both *Drosophila* and human cells. Human patients with loss-of-function mutations in *HBS1L* display several developmental defects, including facial dysmorphia, restricted growth, and hyperpigmented deposits in their retina, and phototransduction defects as measured by electroretinography (ERG) (O’Connell et al. 2019; Luo et al. 2024). Recent work with mouse models has revealed that *HBS1L* deletion results in the loss of several retinal neurons, which likely underlie the vision defects seen in human patients (Luo et al. 2024). However, it remains unknown which cell types in the visual system rely on HBS1L for their proper development and function. Further, specific HBS1L mRNA targets that cause the vision defects have not been described. To address these open questions, we utilized *Drosophila Hbs1* mutants to model the vision defects observed in HBS1L deficiency patients.

We first used ERG measurements to examine whether *Hbs1* mutant animals showed vision defects in comparison to the control wildtype *w^1118^*animals. The compound eye in *Drosophila* is composed of ∼800 individual unit eyes called ommatidia, each containing eight photoreceptors (Sanes and Zipursky 2010). Outer photoreceptors (R1–R6) in the retina project to the first optic neuropil, the lamina, where they synapse with lamina neurons in modular cartridges. A typical ERG trace in control animals contains an initial sustained receptor potential from photoreceptor depolarization, as well as “on” and “off” transients at light onset and offset that reflect synaptic transmission to downstream lamina neurons (*Drosophila* equivalent of bipolar cells) (Heisenberg 1971; Alawi and Pak 1971; Coombe and Heisenberg 1986; Rhodes-Mordov et al. 2015) (**Fig. 3A**). Since ERG analyses are known to be sensitive to eye pigmentation (Stark 1973), we ensured that all our comparative analyses were across animals had similar genetic expression of pigment genes.

**Fig. 3.**
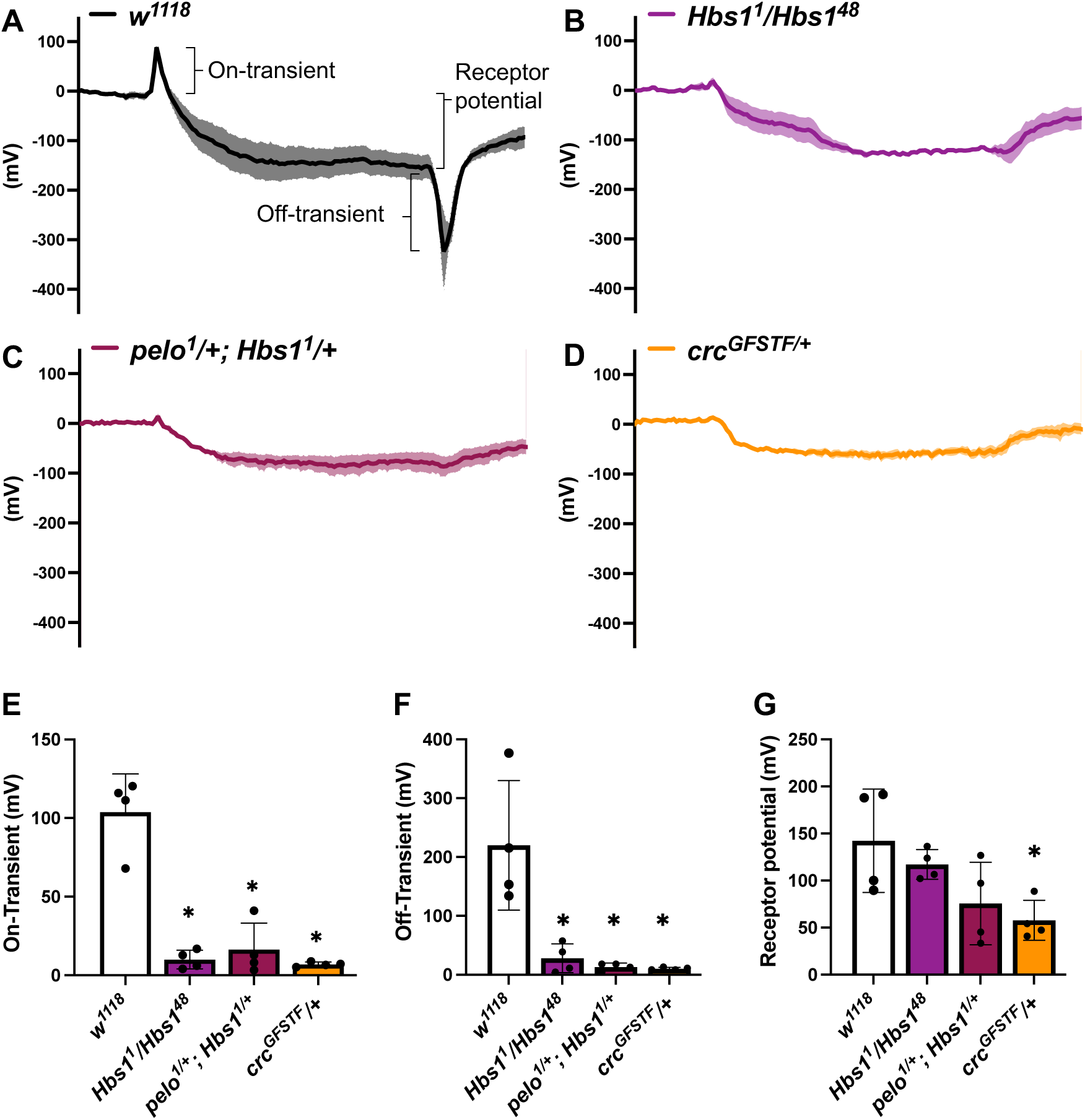
Loss of *Hbs1, pelo,* or *ATF4* results in phototransduction defects. **A-D.** Representative ERG traces from 3-5 day old wildtype (*w^1118^*, A), *Hbs1* mutants (*Hbs1^1^/Hbs1^48^*, B), *Hbs1* and *pelo* hemizygous mutants (*pelo^1^/+;Hbs1^48^/+*, C), and *ATF4* heterozygous mutants (*crc^GFSTF^/+*, D). The solid line represents average of three measurements from an individual animal, and shaded error envelope represents the standard error of mean across the measurements. The three quantifiable parameters of ERGs, on-transient, off-transient, and receptor potential are marked by parentheses in (A). **E-G.** Quantification of on-transient (E), off-transient (F), and receptor potential (G) from (A-D). Data bars are color coded to genotypes in (A-D). Each data point is the average of three independent measurements from a single animal. Data represent the mean from at least four independent animals recorded on separate days; error bars are standard error of mean. Statistical comparisons were made using the Kruskal-Wallis test with Dunn’s correction.

In young (3-5 day old) animals, we found that loss of *Hbs1* resulted in a blunted ERG response to a single light stimulus (**Fig. 3A, B**). Quantification of the ERG parameters showed that *Hbs1* mutants had a significant decrease in on- and off-transients in comparison to control *w^1118^* animals (**Fig. 3E-F**). These data suggested that the vision defects in *Hbs1* mutants may be due to either improper synaptic transmission between the photoreceptor and lamina layers or due to defective lamina function. We also observed a small but statistically insignificant decrease in receptor potential in mutant animals, indicating that photoreceptor depolarization was largely unaffected (**Fig. 3G**). We observed similar results with animals hemizygous for both *Hbs1* and *pelo* (**Fig. 3C,E-G**), suggesting that Hbs1 acts as a complex with Pelo in the context of visual system function. We note here that we were unable to recover homozygous or transheterozygous *pelo^1^ or pelo^PB60^* adults isogenized to the *w^1118^* background and hence opted to test animals hemizygous for *Hbs1* and *pelo*.

Finally, given our results that ATF4 is regulated by Hbs1-Pelo in the fat body (**Fig. 1**), we asked whether loss of ATF4 phenocopies ERG defects seen in *Hbs1* mutants. Since ATF4 (encoded by *cryptocephal, crc* in *Drosophila)* null mutants do not survive to adulthood, we used a partial ATF4 loss of function mutant (*crc^GFSTF^/+*) (Vasudevan et al. 2022). Consistent with our previous report that loss of ATF4 results in age-dependent retinal degeneration, *crc^GFSTF^/+* animals showed blunted ERG traces and reduced on- and off-transients similar to *Hbs1* and *pelo* mutant animals (**Fig. 3D-F**). However, loss of ATF4 also resulted in reduced receptor potential, indicating that these animals may also have defective photoreceptor function in addition to lamina defects (**Fig. 3G**). Nonetheless, these phenotypic analyses demonstrate that loss of Hbs1, its binding partner Pelo, and its mRNA target *ATF4* all result in strikingly similar phototransduction defects.

### Hbs1-Pelo is required in the lamina layer for proper phototransduction

We next sought to use the *Drosophila* genetic toolkit to determine which cell types in the visual system rely on Hbs1-Pelo for proper function. To do so, we used cell type-specific drivers to RNAi deplete *Hbs1* throughout development and subjected young adult animals to ERG analysis. *Hbs1*, *pelo*, and *ATF4* mutants show significant loss in on- and off-transients, which are indicative of lamina defects (**Fig. 3**). Thus, we asked whether depleting *Hbs1* in the lamina using *Gcm-GAL4* (Jenett et al. 2012; Chen et al. 2016) during development was sufficient to recapitulate the ERG defects in *Hbs1* mutants. We observed that 3-5 day old *Gcm>Hbs1^RNAi^* animals showed reduced on- and off-transients with no change to receptor potential (**Fig. 4A, C-E**), consistent with the ERG profiles we observed for *Hbs1* mutants.

**Fig. 4.**
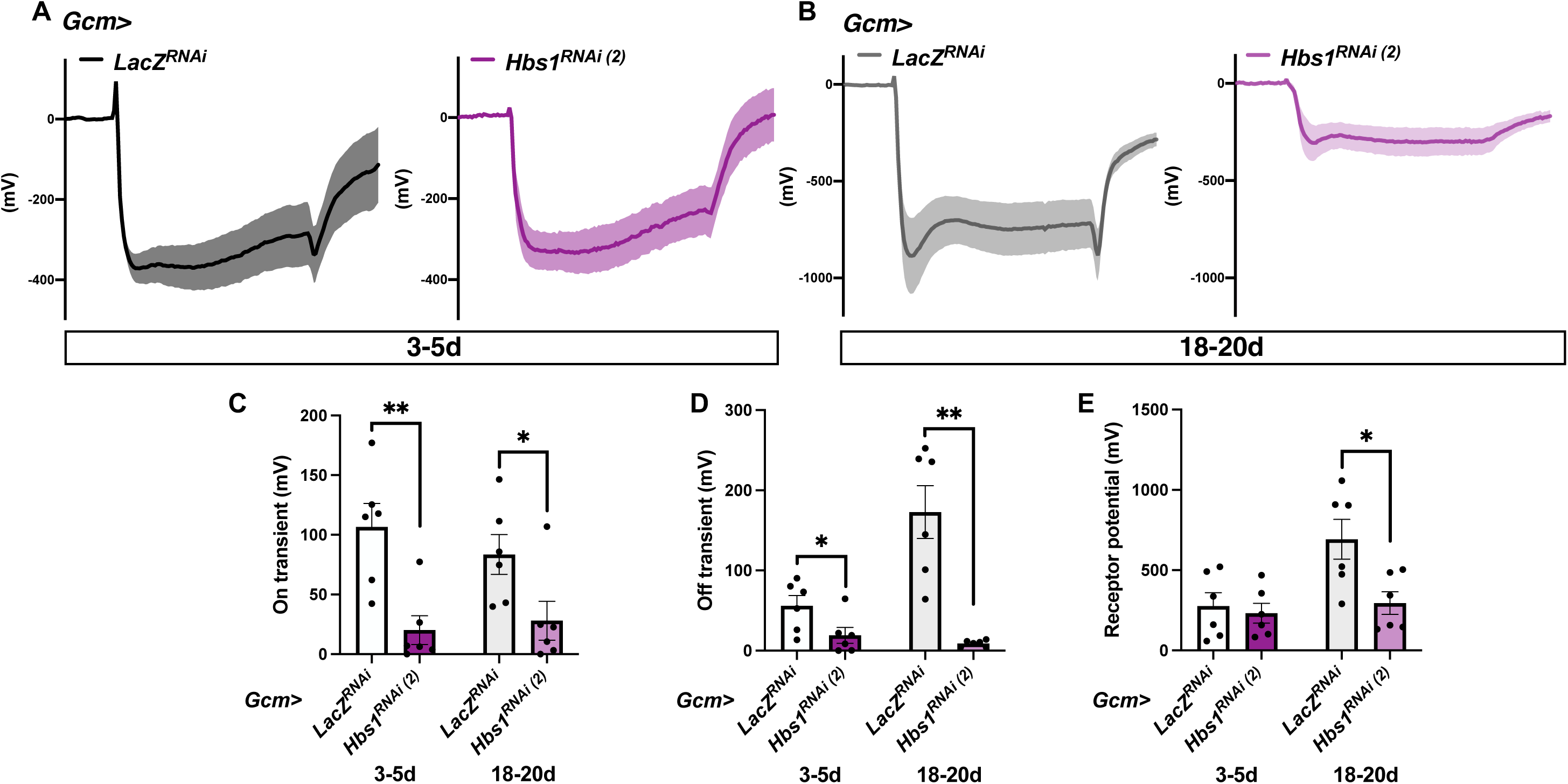
Depleting *Hbs1* in the lamina recapitulates phototransduction defects seen in *Hbs1* mutants. **A-B** Representative ERG traces from 3-5 day old (A) or 18-20 day old (B) animals where a lamina driver (*Gcm-GAL4*) is used to deplete *Hbs1.* Data are graphed as in 3A-D. **C-E.** Quantification of on-transient (C), off-transient (D), and receptor potential (E) from animals in (E-F). Data are plotted as in 3E-G. Statistical comparisons were made using the Mann-Whitney test

The lamina contains five main classes of neurons (L1–L5) that receive direct synaptic input from the outer photoreceptor terminals (R1–R6) (Sanes and Zipursky 2010). It also houses several types of glial cells—epithelial, marginal, and satellite glia—that support lamina neuron function and metabolism. Notably, the *Gcm-GAL4* driver is expressed in epithelial and marginal glial populations and in the lamina neurons. We used a pan-glial driver, *Repo-GAL4* (Sepp et al. 2001), to test whether Hbs1 was required in the lamina glial populations. ERG analysis revealed no significant differences in the ERG parameters between control *Repo>LacZ^RNAi^* and *Repo>Hbs1^RNAi^* animals (**Fig. S3**), suggesting that Hbs1 is not required in lamina glia. This data led us to consider that Hbs1 is likely required in lamina neurons.

Previous studies have demonstrated that defects in lamina neurons can result in an age-related loss in photoreceptor function (Soukup et al. 2013; Lee and Sun 2015). Based on this, we considered whether old *Gcm>Hbs1^RNAi^*animals may exhibit receptor potential defects in ERG analyses, in addition to on- and off-transient defects seen in young animals. Indeed, we found that in comparison to age-matched control *Gcm>LacZ^RNAi^* animals, 18-20 day *Gcm>Hbs1^RNAi^* animals show a marked decrease in their receptor potential, which is reflective of photoreceptor defects (**Fig. 4B, E**). Additionally, we also observed a larger decrease in on- and off-transients in 18-20 day *Gcm>Hbs1^RNAi^* animals when compared to 3-5 day animals (**Fig. 4C-D**), suggestive of age-related further impairment of lamina neuron function as well. Given than *Gcm>Hbs1^RNAi^*produced defects in on- and off-transients in young adults, these data suggest that defects in lamina neuron function may be acquired during development. We also recorded similar ERG defects in *Gcm>pelo^RNAi^* animals (**Fig. 5A-D**), suggesting that Hbs1 acts together with Pelo in the visual system. This led us to propose a model where Hbs1-Pelo is required for lamina neuron development and function, which can in turn impact photoreceptor function with age.

**Fig. 5.**
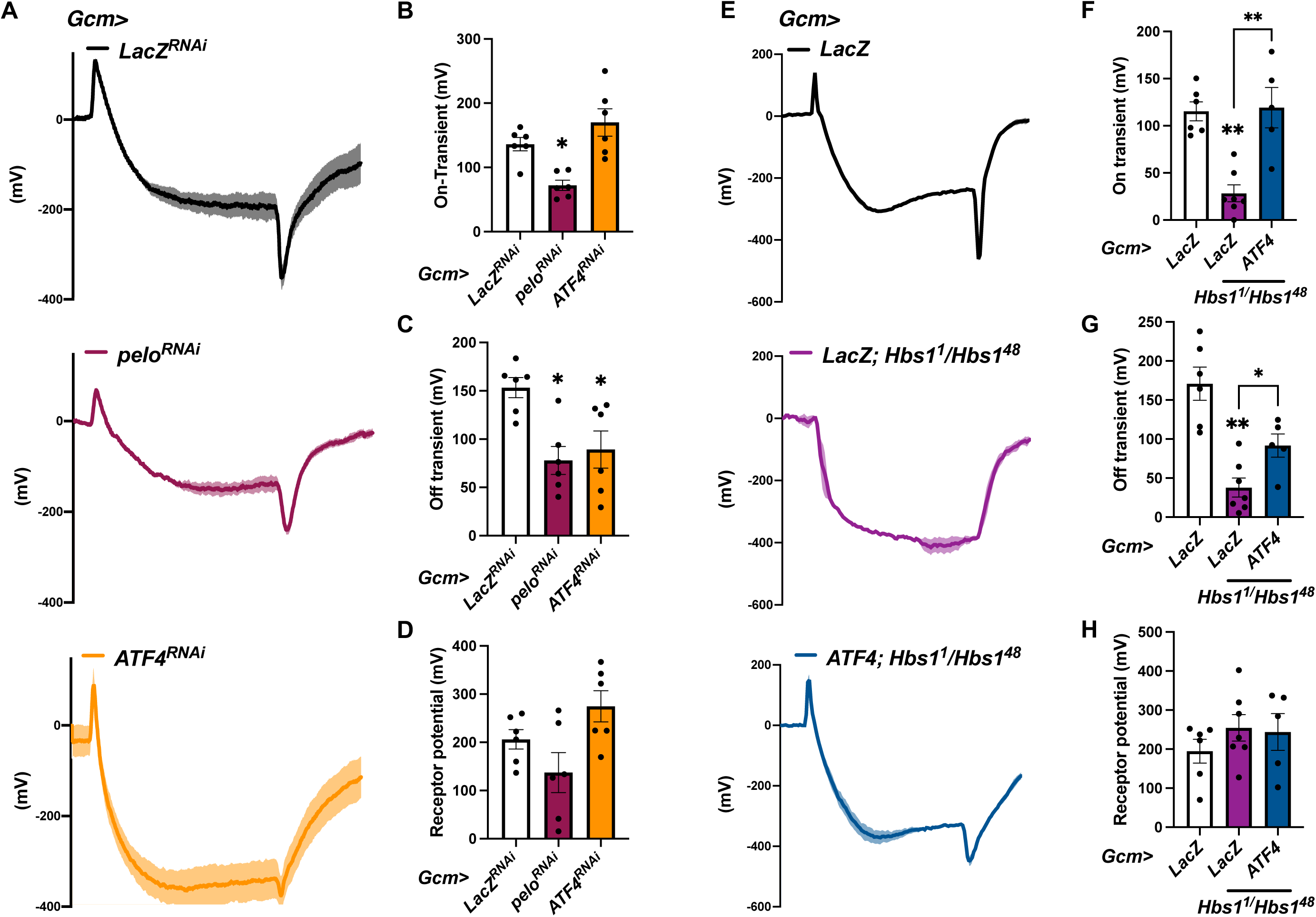
Restoring ATF4 expression in the lamina partially rescues ERG defects in *Hbs1* mutants. **A.** Representative ERG traces from 3-5 day old animals where *Gcm-GAL4* is used to deplete *pelo* (middle, maroon trace), *ATF4* (bottom, orange trace) or express a control *LacZ^RNAi^* (top, black trace). Data are graphed as in 3A-D. **B-D.** Quantification of on-transient (B), off-transient (C), and receptor potential (D) from animals in (A). Data are plotted as in 3E-G. Statistical comparisons were made using Kruskal-Wallis test with Dunn’s correction. **E.** Representative ERG traces from 3-5 day old animals where *Gcm-GAL4* drives expression of a control *UAS-LacZ* transgene in wildtype animals (top, black trace), *Hbs1* mutants (middle, purple trace) or of a leaderless *ATF4* in *Hbs1* mutants (bottom, blue trace). Data are graphed as in 3A-D. **F-H.** Quantification of on-transient (F), off-transient (G), and receptor potential (H) from animals in (A). Data are plotted as in 3E-G. Statistical comparisons with *Gcm>LacZ* were made using the Kruskal-Wallis test with Dunn’s correction and are indicated as floating asterisks when significant. Other comparisons were made using the Mann-Whitney test.

### Restoring ATF4 in the lamina partially rescues phototransduction defects in Hbs1 mutants

Though previous translation profiling studies in *HBS1L* deletion patient fibroblasts showed changes in translation efficiency of many mRNAs (O’Connell et al. 2019; Luo et al. 2024), none of them have been linked to the etiology of vision defects observed in patients. Given our findings that HBS1L-Pelo regulates ATF4 (**Fig. 1, 2**) and the phenotypic similarities in ERGs from *Hbs1* and *ATF4* mutants (**Fig. 3**), we asked whether the vision defects with loss of *Hbs1* may partly be due to reduced ATF4 expression. To test this possibility, we first examined whether depleting *ATF4* in the lamina phenocopies phototransduction defects seen with lamina-specific *Hbs1* depletion. ERG measurements from young *Gcm>ATF4^RNAi^*animals showed decreased off-transients but with no change to on-transient or photoreceptor potential (**Fig. 5A-D**), suggesting a possible role for ATF4 in the lamina.

We next asked whether restoring ATF4 expression in the lamina was sufficient to rescue the ERG defects seen in *Hbs1* mutants. To do so, we drove expression of a leaderless *UAS-ATF4* (Vasudevan et al. 2020) or a control *UAS-LacZ* using *Gcm-GAL4* in *Hbs1^1^/Hbs1^48^* animals. Consistent with our data in *Hbs1* mutants (**Fig. 3A-B, E-G**), our ERG analyses revealed that *Gcm>LacZ* expression resulted in on- and off-transient defects in *Hbs1^1^/Hbs1^48^* animals when compared to control *w^1118^* animals (**Fig. 5E-G**). These defects in *Hbs1^1^/Hbs1^48^* animals were partially rescued by *Gcm>ATF4* expression (**Fig. 5E-G**), demonstrating that ATF4 is a relevant downstream target of Hbs1 in the lamina. These genetic analyses together suggest that Hbs1-Pelo mediated regulation of ATF4 is required for proper lamina function.

### *Hbs1* mutants show vacuolation defects in the lamina layer

Since our phenotypic analyses with ERG revealed that vision defects in *Hbs1* mutants stem from improper lamina neuron function, we used confocal microscopy to investigate the lamina cortex and neuropil in *Hbs1^1^/Hbs1^48^* animals. We stained whole eye-optic lobe preparations from *Hbs1* mutants with a pan-neuronal marker ELAV (Robinow and White 1988; 1991). Z-stack analysis of confocal images revealed that loss of *Hbs1* had no obvious impact on organization of the lamina neuron cell bodies in comparison to wildtype animals (**Fig. 6A-B**). We further probed these results with lamina neuron subtype markers (Chen et al. 2010; Hasegawa et al. 2013; Fernandes et al. 2017) to examine whether loss of *Hbs1* impacted only a subpopulation of L1-L5 neurons which may be obscured by using a pan-neuronal marker. However, these analyses did not reveal any differences in the number of individual L1-L5 subtypes or total number of lamina neurons between *w^1118^* wildtype and *Hbs1* mutants (**Fig. 6C, S4**). We also used a pan-glial marker, Repo (Xiong et al. 1994; Halter et al. 1995), and observed no differences between wildtype and *Hbs1* mutants in either the arrangement or numbers of epithelial and marginal glia, which can be distinguished based on their location in the lamina neuropil (**Fig. 6A-B**, **D-E**). These data led us to consider that the ERG defects in *Hbs1* mutants are unlikely to be caused by disruptions to lamina neuron or glial specification or survival, but instead may arise due to defects in synaptic transmission between the photoreceptors and lamina neurons.

**Fig. 6.**
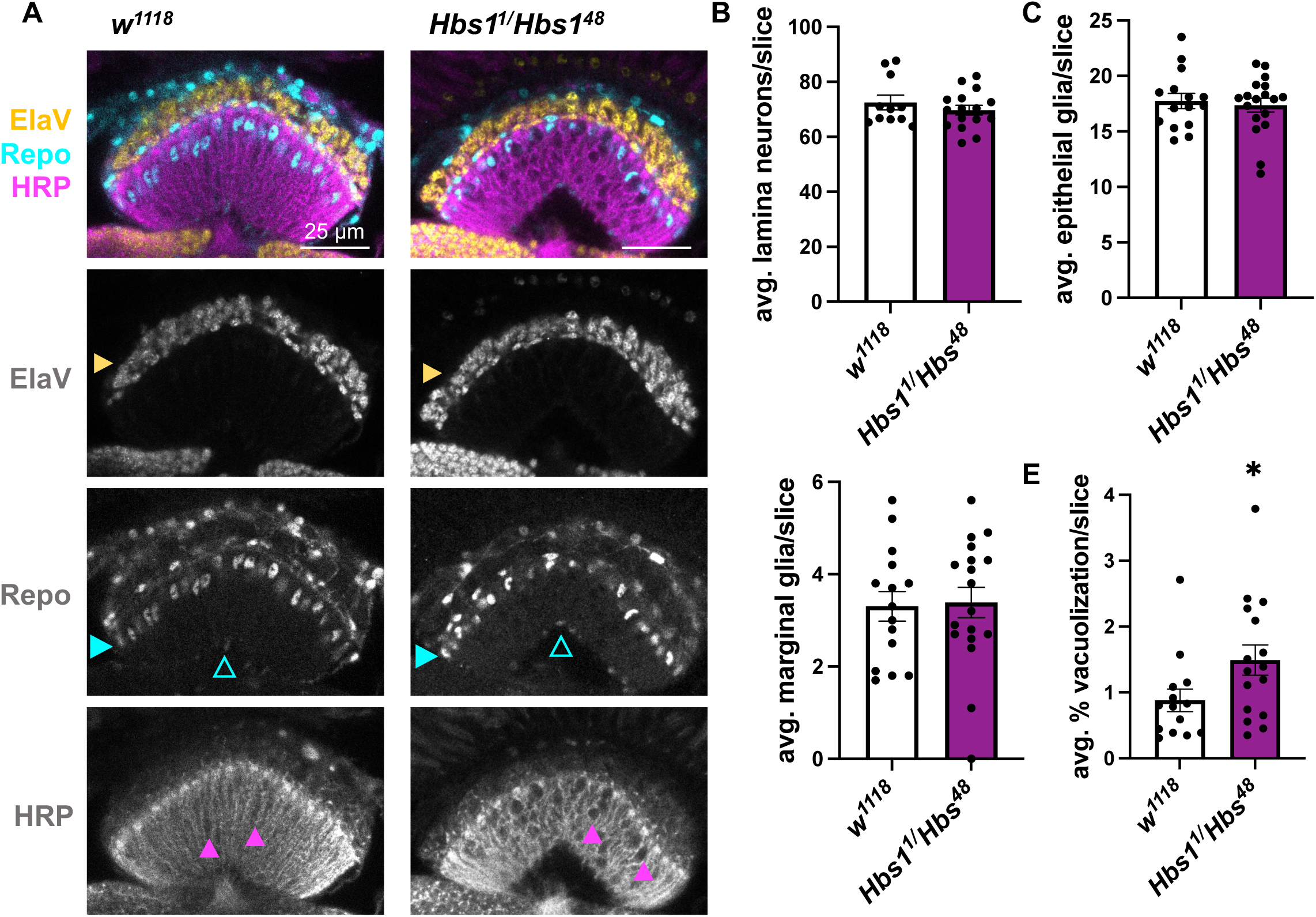
Loss of *Hbs1* results in increased vacuolization in the lamina neuropil with no effect on the number or morphology of lamina neurons and glia. **A.** Confocal images of 3-5 day old adult lamina from wildtype or *Hbs1* mutant animals stained with a neuronal soma marker (Elav, yellow), glia soma marker (Repo, cyan), and neuronal membrane marker (HRP, magenta). Yellow solid arrows point to lamina neurons, cyan solid arrows point to epithelial glia, cyan open arrows point to marginal glia, magenta solid arrows point to areas of vacuolization. **B.** Quantification of the total number of lamina neurons in wildtype and *Hbs1* mutants as calculated by summation of individual lamina neurons subtypes (L1-L5) from S4A-B. Each data point represents one animal, data bars represent the mean of at least ten animals, error bars represent standard error of mean. **C-D.** Quantification of the number of epithelial (C) and marginal (D) glia from (A). Epithelial glia are identified as Repo-positive cells distal to the lamina neuropil and marginal glia are Repo-positive cells proximal to the lamina neuropil (see cyan arrows in A). Each data point represents one animal, data bars represent the mean of at least ten animals, error bars represent standard error of mean. **E.** Quantification of the percentage of vacuolization as measured by the area of the vacuoles normalized to the total area of the lamina neuropil. Each data point represents one animal, data bars represent the mean of at least ten animals, error bars represent standard error of mean. Statistical comparisons were made using the Mann-Whitney test.

To test for synaptic disruptions in the lamina neuropil, we stained eye-optic lobe preparations with the neuronal membrane marker, HRP (Jan and Jan 1982). Strikingly, HRP staining in *Hbs1* mutants revealed an increased incidence of vacuoles within the neuropil, visible as dark ‘holes’ in the neuropil (**Fig. 6A-B, F**). Vacuolization can arise from defects in synapse formation/maintenance, photoreceptor axon transport, or glial support (Coombe and Heisenberg 1986; Jackson et al. 2002; Iijima-Ando et al. 2012; Lee and Sun 2015). However, our ERG analyses do not suggest defects in photoreceptors based on unchanged receptor potential in young animals (**Fig. 3**), and depleting *Hbs1* in glia did not result in ERG defects (**Fig. S3**). Based on these eliminatory analyses and considering all our combined data, we conclude that Hbs1-Pelo likely regulates ATF4 expression in lamina neurons and that such regulation is required for proper synaptic transmission between the lamina and photoreceptor layer.

## Discussion

Inherited retinal diseases are a varied group of genetic disorders that cause progressive vision loss due to gradual photoreceptor death (Duncan et al. 2024). While most of the over 260 genes identified for these diseases are specifically expressed in retinal cells, more broadly expressed ribosome-associated proteins have also been associated with retinal degeneration. Specifically, mutations in RPL10, GTPBP1, GTPBP2, and HBS1L have been shown to result in progressive vision loss, in addition to general neurodegeneration (Zanni et al. 2015; Ishimura et al. 2014; O’Connell et al. 2019; Terrey et al. 2020), with much ongoing research aimed at understanding the molecular underpinnings of these diseases. Our work here uses a *Drosophila* model to significantly advance our etiological understanding of vision defects associated with HBS1L deficiency and provides a preliminary molecular mechanism for progressive vision loss in patients.

### Regulation of translation reinitiation by termination factors

Our investigation of vision defects in *Hbs1* mutants were rooted in our efforts towards understanding regulation of the ISR transcription factor, ATF4. The expression of ATF4 is tightly regulated at multiple stages, including transcription, transcript stability, translation, and protein stability. Among these, regulation of mRNA translation within the ATF4 5’ leader is the best characterized and arguably, the most impactful on ATF4 protein expression. The human ATF4 5’ leader has two uORFs, with the final uORF2 overlapping with the ATF4 ORF. A substantial body of work has investigated the signaling events that favor delayed reinitiation at the ATF4 ORF in lieu of reinitiation at uORF2, predominantly the regulation of initiator methionine availability by the ISR pathway. Regardless, it remains that there is at least one, if not two, translation termination events (at the start-stop and at uORF1) within the 5’ leader that precedes reinitiation at the ATF4 start codon. Our data implies a role for efficient ribosome splitting by HBS1L-Pelo at these upstream stop codons (**Fig. 2D**), which is a necessary precursor to reinitiation at the ATF4 start codon.

High-resolution ribosome profiling data from the Teleman laboratory has convincingly demonstrated that DENR-MCTS1 or eIF2D is required to evacuate spent tRNAs from codons penultimate to the stop codon and thus promote faithful termination (Bohlen et al. 2020). Their meta-analyses further revealed that these factors were specifically required at certain codon identities, including GCG^Ala^ which is the penultimate codon in the uORF1 of the ATF4 mRNA across many species. Such bias towards codon identity for the action of DENR-MCTS1/eIF2D likely stems from their preferential binding to specific tRNAs. Based on the molecular function of HBS1L-Pelo, it is unlikely that penultimate codon identity is relevant to their action on the ATF4 5’ leader. Notably, another Pelo was implicated in translation reinitiation of another stress responsive uORF-containing transcript, C/EBPα (Fernandez et al. 2024). It remains to be investigated whether the presence of uORFs is sufficient for an mRNA to be targeted by HBS1L-Pelo, or whether there are other sequence features in an mRNA that render such selectivity.

Previous mechanistic studies have demonstrated that HBS1L-Pelo promote ribosome recycling on mRNAs that are targeted for no-go decay and no-stop decay, which are mRNA surveillance mechanisms that identify and degrade faulty mRNA transcripts (Shoemaker et al. 2010; Saito et al. 2013). Intriguingly, the *ATF4* mRNA is a known target for another mRNA surveillance mechanism, nonsense-mediated decay (NMD) target (Wang et al. 2011; Wengrod et al. 2013) raising the question of whether the effects of HBS1L-Pelo on ATF4 are linked to its role in NMD. However, our data shows that depleting HBS1L or Pelo does not impact *ATF4* mRNA levels (**Fig. S2A**), which would preclude the involvement of mRNA degradation mechanisms such as NMD.

### Selective mRNA translation in the visual system

An increasing body of literature provides evidence that translational regulation drives visual system development to a similar extent as classic transcriptional programs (Jung and Holt 2011; Zhang et al. 2016; Chen et al. 2021; Ichinose et al. 2024) and others). Under this paradigm, the presence of a mRNA does not necessarily imply that the corresponding protein is translated.

Instead, cellular mRNAs are selectively translated due to their sequence features. We propose that in some cases, this selectivity falls to specialized translation factors such as EIF3H, DENR, MCTS1, EIF2D as exemplified by their role in regulating translation reinitiation in the ATF4 5’ leader (Roy et al. 2010; Bohlen et al. 2020; Vasudevan et al. 2020). Our work herein extends this list to HBS1L and Pelo.

Our data showed that *ATF4* mutants exhibit receptor potential defects (**Fig. 3G**), which we do not observe with *Hbs1* or *pelo* loss of function, implying that other factors are likely required for the proper expression of ATF4 in photoreceptors. Likewise, specialized factors such as HBS1L-Pelo likely have other relevant mRNA targets, both in the visual system and elsewhere. For instance, *Drosophila Hbs1* and *pelo* mutants display defective spermatogenesis, though such a phenotype has not been reported in *ATF4* mutants and it remains unknown which mRNAs downstream of Hbs1-Pelo effect this phenotype. The best way to identify the targets for these specialized factors is through cell-type-specific ribosome profiling, a technically challenging method that has recently yielded promising results (Ichinose et al. 2024). High-resolution translational profiling combined with comparative analyses using model organisms is expected to reveal more such paradigms in the future.

### ISR signaling in visual system development and function

Recent research from multiple groups has implicated the ISR pathway in the visual system, particularly in the context of retinal disorders like autosomal dominant retinitis pigmentosa (adRP) (Bhootada et al. 2016; Athanasiou et al. 2017; Comitato et al. 2019; Vasudevan et al. 2020; 2022; Zhao et al. 2023). Nearly 25% of adRP cases are caused by misfolding-prone mutations in Rhodopsin (Lewin et al. 2014). The prevailing model suggests that misfolded rhodopsin activates the ER stress-sensitive ISR kinase, PERK, which initiates a protective transcriptional program. Predictably, loss of *PERK* exacerbates retinal degeneration in a *Drosophila* adRP model (Vasudevan et al. 2020). The protective function of PERK in photoreceptors is attributed this cell type’s high secretory load of opsins, which makes it particularly reliant on maintaining ER homeostasis (Zhang et al. 2014), amongst other causes. Such a protective role has been extended to other factors downstream of PERK, including ATF4 and its regulator, eIF2D, and also other ER stress response pathways (Vasudevan et al. 2022; 2020; Yan et al. 2019; Coelho et al. 2013; Ryoo et al. 2007). Based on this, we predict that loss of *Hbs1* and *pelo* will likely also result in exacerbated retinal degeneration in the *Drosophila* adRP model.

Our findings here using ERG analyses in *ATF4* mutants (**Fig. 3**) extends the role of ISR signaling beyond the context of adRP, to visual system development and function. We previously demonstrated that partial loss-of-function mutants of *ATF4* exhibit age-dependent retinal degeneration (Vasudevan et al. 2022). Others have confirmed expression of an ATF4 5’ leader reporter in multiple cell types within the *Drosophila* visual system, including photoreceptors (Kang et al. 2015). In photoreceptors, the role for PERK-ATF4 and other pathways involved in maintaining ER homeostasis appears intuitive due to the highly secretory nature of this cell type. Consistently, our data also shows that loss of ATF4 function results in a decreased receptor potential in young flies (**Fig. 3G**). However, our data also demonstrate a crucial and previously undescribed role for ATF4 in lamina neurons, as evidenced by 1) diminished on- and off-transients in *ATF4* loss of function mutants (**Fig. 3E-H**), and 2) the rescue of on- and off-transients in *Hbs1* mutants by restoring ATF4 expression (**Fig. 4E-H**). Whether and how ISR signaling is activated in these second order neurons during development is an open question, and the molecular role of ATF4 in synapse formation between the lamina and photoreceptors is less intuitive. These avenues bear clear pharmacological interest and warrant further investigation.

### Implications of lamina Hbs1 function in human HBS1L deficiency-related vision loss

Our data clearly demonstrate a role for Hbs1 in the lamina, as seen by ERG (**Fig. 4**) and confocal analyses (**Fig. 6**). We further narrowed down Hbs1 function to lamina neurons, since depleting Hbs1 in glia using *Repo-GAL4* did not show any ERG defects (**Fig. S3**). It is possible that there are other cell types that require Hbs1-Pelo for their function that our study did not capture. Despite these other potential roles, the involvement of Hbs1 in lamina neurons is made evident by the observed vacuolization defects (**Fig. 6**).

Both *Drosophila* lamina neurons and human bipolar cells act as interneurons that are responsible for relaying and processing visual information from photoreceptor cells (Sanes and Zipursky 2010; Malin and Desplan 2021). In the fly, the R1-R6 photoreceptors terminate in the lamina and synapse with L1-L5 lamina neurons, which then transmit this information to the medulla. Similarly, in the human eye, photoreceptors (rods and cones) synapse with bipolar cells, which then relay the signal to the retinal ganglion cells. Our extensive genetic analyses implicate *Drosophila* Hbs1 and its binding partner, Pelo, in proper function of the lamina neurons (**Fig. 3-4**), specifically in synapse formation between the lamina neurons and photoreceptors (**Fig. 5**). Extrapolating these analyses to the human visual system, we propose that HBS1L is likely required for proper development of bipolar neurons. There is no direct evidence in other models that loss of Pelo results in vision defects. However, previous work has shown that human HBS1L deficiency patient fibroblasts show lower levels of Pelo protein (O’Connell et al. 2019), suggesting that at least some cell types in HBS1L deficiency patients also have reduced Pelo. This further supports a model wherein HBS1L acts together with Pelo in visual system development and function.

As with the *Drosophila* ERG, the human ERG response can be quantified by the amplitude and timing of specific changes in electrical potential, called the a-wave and b-wave (Gauvin et al. 2018). The a-wave represents the electrical activity of the photoreceptors (rods and cones), indicating their health and function. The subsequent b-wave reflects the activity of the bipolar cells and Müller glia, which are responsible for transmitting the signal from the photoreceptors to the inner retina. Consistent with our conclusion that HBS1L is required primarily in bipolar cells, ERG readings from human HBS1L deficiency patients have been reported to show dampened b-wave amplitudes (Luo et al. 2024). However, human ERG readings from HBS1L deficiency patients also show reduced a-wave amplitudes, indicative of defective photoreceptor (particularly cone cell) function (Luo et al. 2024), which is reminiscent of reduced receptor potential in *Hbs1* mutants with age (**Fig. 4I**). Thus, we consider that the cone cell impairment in HBS1L deficiency patients is an age-dependent secondary consequence of improper synapse development between photoreceptors and bipolar neurons. Indeed such a mechanism has been demonstrated in some mouse models of retinal degeneration disorders (Ou et al. 2015; kleine Holthaus et al. 2018) and observed with lamina neuron defects in *Drosophila* (Soukup et al. 2013; Lee and Sun 2015). This hypothesis is also somewhat supported by retina images from mouse *HBS1L* deletion mutants, which showed fewer photoreceptors at 14 days postnatal but not 7 days postnatal (Luo et al.

2024). Further, whole retina images from *HBS1L* mutant mice show reduced thickness in the outer plexiform layer (Luo et al. 2024), which houses the synapses between photoreceptors and bipolar cells and is the equivalent of the lamina neuropil (Sanes and Zipursky 2010; Malin and Desplan 2021). Finally, transcriptomic analyses from the mouse retina show that HBS1L is expressed in bipolar cells and also in photoreceptor cells (Luo et al. 2024). Taking the data across these multiple models together, we speculate that HBS1L-Pelo is required in bipolar cells for proper synapse formation with photoreceptors. Thus, our work herein proffers the first known mechanistic framework for understanding the basis of vision defects in HBS1L deficiency patients.

## Methods

### *Drosophila* husbandry and stocks

All stocks and crosses were reared at 25°C with a 12-hour light/dark cycle. Progeny of interest were collected within 2 days of eclosion and aged in vials with no more than 10 animals each aged at 25°C. Aging animals were transferred to new vials every 3 days to ensure fresh food availability. Animals were raised on ‘R’ food formulation from Lab Express (https://www.lab-express.com/DIS58.pdf). All fly stocks used in the study, and the specific figures wherein they are employed are listed in **Table S2**.

Sex as a biological variable: Initial observations were recorded in both males and females to ensure that the phenotypes were not sexually dimorphic. All the final quantified analyses performed in this study utilize male animals in both larval and adult experiments for experimental consistency.

### Immunostaining and confocal microscopy

Fat bodies from wandering third instar male larva were dissected in 1X phosphate buffer saline (PBS) and placed in an Eppendorf tube. Tissues were fixed in 4% paraformaldehyde (PFA), 1X PBS for 20 minutes on a nutator, and washed twice for 10 minutes each in 0.1% PBS-Tween (PBST). Samples were incubated for 10 minutes in the dark with 4′,6-diamidino-2-phenylindole (DAPI, 200 μM final) to counterstain for nuclei. Fat bodies were finally mounted on glass slides in a solution of 70% glycerol prior to placing coverslips. Slides were sealed with a thin coating of nail polish to prevent evaporation and imaged on a Nikon A1 inverted line scanning confocal microscope.

Adult whole brains (optic lobes and central brain) were dissected in 1X PBS and placed in a glass dish, where they were fixed in 4% PFA for 30 minutes. Samples were washed with 0.5% PBTx (1X PBS with 0.5% TritonX) for at least 1 hour, and then incubated with primary antibodies diluted in block solution (5% normal horse serum in PBTx) for 48 hours at 4°C. After washing with 0.5% PBTx, samples were further incubated for 48 hours at 4°C with secondary antibodies diluted in block, washed again, and finally mounted in SlowFade (Life Technologies). Slides were imaged on a Leica SP8 upright point scanning confocal, with 40× objective and 2x zoom, and stacks were acquired with a step size of 1 μm.

Antibodies: Rat anti-Elav (DSHB 7E8A10, 1:20), mouse anti-Repo (DSHB 8D12, 1:20), DyLight 405 conjugated goat anti-Horseradish Peroxidase (HRP) (Jackson ImmunoResearch 123-475-021, 1:50), guinea pig anti-Slp2 (a gift from C. Desplan, 1:200), mouse anti-Svp (DSHB 2D3, 1:20), rat anti-Erm (a gift from C. Desplan, 1:100), rabbit anti-Bsh (a gift from C. Desplan, 1:400). Secondary antibodies were used at 1:400 for Alexa Rhodamine Red-X anti-rat (Jackson ImmunoResearch, 712-295-153), Alexa Fluor 647 anti-mouse (Jackson ImmunoResearch, 715-605-151), Alexa Fluor 488 anti-quinea pig (Jackson ImmunoResearch, 706-545-148), Alexa Fluor Rhodamine Red-X anti-mouse (Jackson ImmunoResearch, 715-295-151), and Alexa Fluor 647 anti-rat (Jackson Immunolabs, 712-605-153), and 1:50 for Alexa Fluor 405 anti-rabbit (Invitrogen, A48258).

#### Image quantification

Fat body: Images were analyzed in ImageJ v2.1.0/1.53c. Using the DAPI channel and the ‘Threshold’ function, individual nuclei were identified as regions of interest (ROI). DsRed intensity was then collected for each ROI and graphed.

Lamina: Images were analyzed in ImageJ v2.1.0/1.53c. The Cell counter plug-in was used for lamina neuron subtype counts. Svp, Slp2 double positive cells were counted as L1 neurons. Slp2 single positive cells were counted as L2 neurons. Erm positive cells were counted as L3 neurons. Bsh positive cells were counted as L4 neurons, and Slp2, Bsh double positive cells were counted as L5 neurons. The number of cells were counted in each z-stack slice and averaged across equal number of slices for every sample to create a single data point for each sample. The sum of the lamina neuron subtypes were calculated for the total lamina neuron counts. Epithelial glial counts were done with ImageJ cell counter plug-in and were identified as Repo-positive cells distal to the lamina neuropil, while marginal glia are Repo-positive cells proximal to the lamina neuropil. Percent vacuolization was quantified by the sum of the areas of each vacuole (regions lacking HRP staining) in one neuropil divided by the total area of the neuropil. Each data point represents the average percent vacuolization across z stacks from one animal.

### Cell culture and western blot analyses

HEK293T cells were cultured in high glucose Dulbecco’s Minimal Essential Medium (DMEM) supplemented with 10% Fetal Bovine Serum and 1% Penicillin/Streptomycin. siRNA and plasmid (catalog details below) co-transfections were performed using Lipofectamine 2000 (Life Technologies) according to manufacturer’s protocol in 6-well dishes. 48hrs after transfection, cells were washed twice in 1x PBS and lysed in 100 μl of Radioimmunoprecipitaton assay (RIPA) buffer (50 mM Tris-HCl pH 7.4, 150 mM NaCl, 1% Triton X-100, 0.5% Sodium deoxylcholate, 0.1% SDS, 1 mM EDTA, protease inhibitor cocktail), and the lysate was cleared by centrifuging for 10 minutes at 14000 rcf at 4°C. 20μl of the lysate was mixed with the appropriate amount of 4x Laemmli buffer (277.8 mM Tris-HCl pH6.8, 40% v/v glycerol, 4% SDS, 0.02% bromophenol blue, 4% ϕ3-mercaptoethanol) and analysed by western blotting on nitrocellulose membrane. Membranes were blocked for 1 hour at room temperature in 5% non-fat dry milk diluted in PBST. They were then incubated in primary antibodies (listed below) diluted in 5% BSA in PBST overnight at 4°C. Following three 10-minute washes in PBST, membranes were incubated in secondary antibodies for 2 hours at room temperature and washed thrice for 10 minutes in PBST before HRP detection.

Puromycin assays: Cells transfected as above were treated with 10 μg/ml for 15 minutes prior to lysis. Lysates were then analyzed by western blotting as described above.

Antibodies: ATF4 (Cell Signaling 11815S, 1:1000 in 5% BSA PBST, HBS1L (Proteintech 10359-1-AP, 1:2000 in 5%BSA PBST), Pelo (Proteintech 0582-1-AP, 1:2000 in 5% BSA PBST), Tubulin (DSHB 12G10,1:2000 PBST), GFP (Immunology consultants 50-196-2090, 1:2000 PBST), Puromycin (Sigma MABE343, 1:2000 in 5% BSA PBST). Host-appropriate HRP-conjugated secondary antibodies (Jackson Immunolabs) were used in all cases except for Tubulin, where a Starbright700-conjugated secondary (Biorad) was used. at All secondary antibodies were used at 1:5000 in PBST.

qPCR: Cells transfected as above were washed twice with 1x PBS and total RNA was collected by adding Trizol (Life Technologies) directly to the cell culture dish following manufacturer’s protocol for RNA preparation. 1000ng of RNA was used to in a 20 μl cDNA reaction using the Thermo Maxima reverse transcriptase (Life Technologies) following manufacturer’s protocol. Quantitative RTPCR analysis was performed using SYBR Green (Mid Sci) and relevant primers listed below.

Nucleic acids: Control siRNA (Thermo 4390843 Negative control siRNA), *HBS1L* siRNA (Thermo Silencer Select s21152), Pelo siRNA(Thermo Silencer Select s28807); ATF4 5’ leader-GFP (Addgene 21852, 5’ATF4.GFP), Mutant ATF4 5’ leader-GFP (Addgene 21863, 5’ATF4.uORF1stopm1&2.GFP)

#### qPCR Primers

ATF4 - F: GGAGATAGGAAGCCAGACTACA, R: GGCTCATACAGATGCCACTATC GAPDH - F:GTCTCCTCTGACTTCAACAGCG, R:ACCACCCTGTTGCTGTAGCCAA

### ERG recordings and analyses

All ERG recordings were performed in an electrophysiology rig covered with a black out Faraday cage following a protocol derived from prior studies (Vilinsky and Johnson 2012; Wu et al. 2022; Meece et al. 2025). *Drosophila* were anesthetized by placing them on ice for 10 minutes and immobilized using dental wax on glass coverslips for electrophysiological recordings as described before (Meece et al. 2025). Reference and recording electrodes were prepared as in (Vilinsky and Johnson 2012) and clamped into standard electrode holder (Warner Instruments E series, straight configuration). The reference electrode was generated using glass capillaries (World Precision Instruments, model 1B150F-4) in vertical pipette puller (David Kopf instruments) to approximate tip size of 0.5μm and filled with 0.9% w/v NaCl solution. The recording electrode is generated by inserting a cotton sewing thread into the barrel of an (un-pulled) glass capillary then filled with 0.9% w/v NaCl. Using two micromanipulators (World Precision Instruments), the reference electrode was inserted in the thorax just below the wing, and the recording electrode was gently placed on the external eye. The recording electrode was connected to the input headstage of an A/M Systems headstage Model 3000 (Regular) amplifier, and the reference electrode was connected to a common aluminum ground bar located inside the Faraday cage. Data from the amplifier were routed to an A/D converter (AD Instruments PowerLab 4/30) and subsequently acquired, analyzed and displayed using the program LabChart 8 (AD Instruments). Animals were stimulated with a 1s exposure to blue LED (wavelength: 470 ± 20 nm, Phillips Luxeon Rebel LED). Each animal was measured three times, with a one-minute rest between recordings. All ERG analyses were performed from animals collected across at least two independent crosses reared in identical conditions, using similar ERG setups.

### Statistical analysis

The figure legends describe the sample sizes for the corresponding data. Multiple comparisons with a control were made using a Kruskal-Wallis test with Dunn’s correction for multiple comparisons. Paired comparisons were made using the Mann-Whitney test for nonparametric data in all instances when we could not assume Gaussian distribution. For data where control samples were normalized to 1 (such as western blots or qPCRs), a Wilcoxon signed rank test was conducted when comparing a given sample the control. These data sets have matched data, where each data point has a corresponding measurement across all conditions, thus when making multiple comparisons we utilized a Friedman test for nonparametric data with Dunn’s correction for multiple comparisons. When comparing two conditions in these data sets, we used a Wilcoxon matched-pairs signed rank.

## Acknowledgements

We would like to thank publicly available model organism resources that fueled our research: FlyBase, Bloomington Drosophila stock center, and Vienna Drosophila stock center. The *Hbs1* mutants (*Hbs1^1^* and *Hbs1^48^*) were a generous gift from Dr. Rongwen Xi. We are grateful to Drs. Ilya Vilinskiy and Atulya Iyengar for their help with our ERG experimental set up and analyses. We would like to thank all members of our lab for discussion and feedback on the project. Stocks obtained from the Bloomington Drosophila Stock Center (NIH P40OD018537) and Vienna Drosophila stock center (Dietzl et al. 2007) were used in this study. We used FlyBase (release FB2024_01) for identifying phenotypes and stocks in this study.

## Contributions

D.V. conceived the project. D.V. and V.M. F. designed the experiments. D.V. wrote the paper. N.N., C.G., and D.V. performed the ERG analyses, and A.C. performed pilot cell culture experiments. I. L-B. performed the adult eye dissections and imaging. K.Q. performed all cell culture experiments and microscopy quantifications.

## Funding

K.Q., N.N., C.G., A.C., and D.V. were supported by NIH R35GM150516 (to D.V.) and NIHR00EY029013 (to D.V.). I.L-B and V.M. F were supported by a Wellcome CDA (225986/Z/22/Z), EMBO Young Investigator Award and Lister Institute Prize (all to V.M.F).

---

**Fig. S1.**
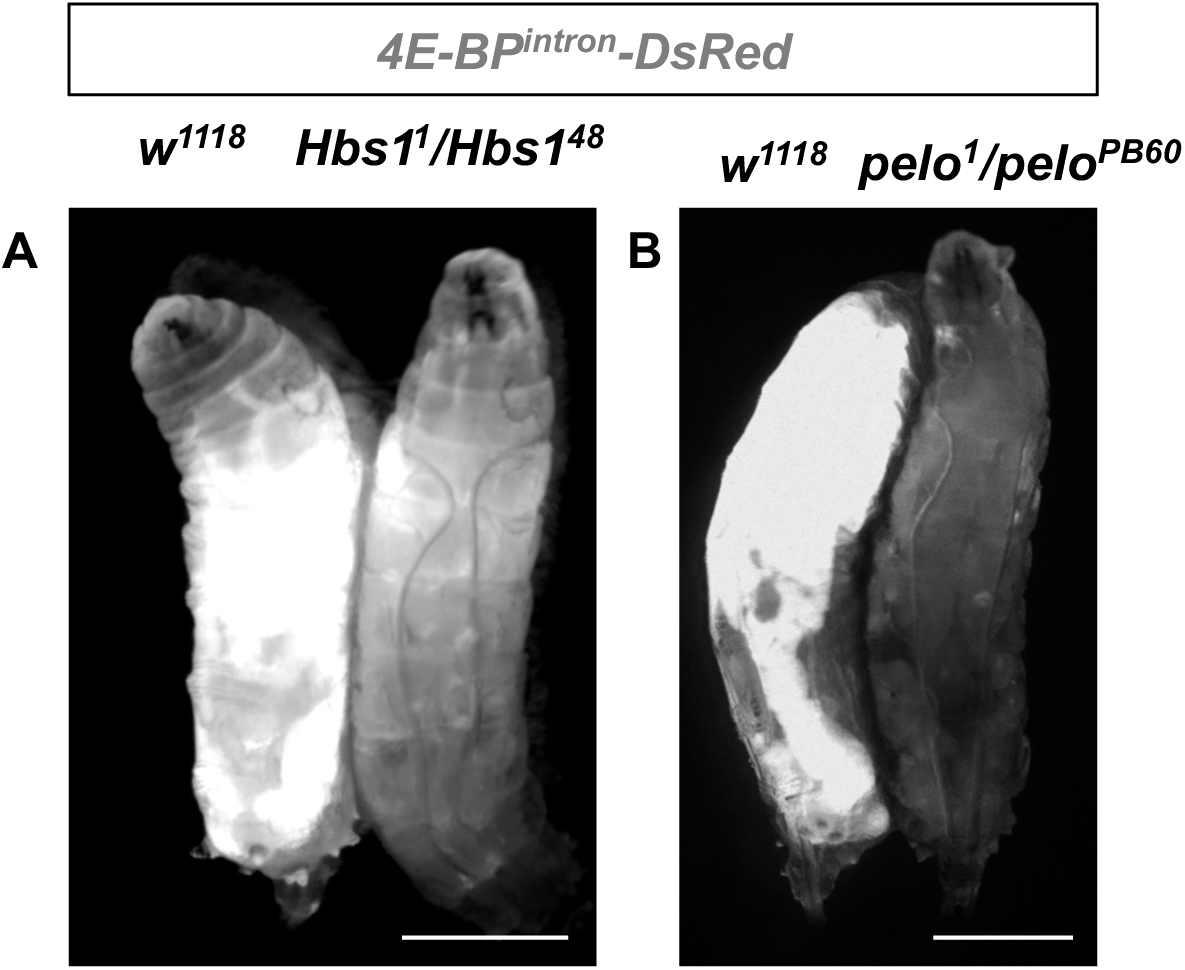
Loss of *Hbs1* or *pelo* results in reduced ATF4 reporter expression. Fluorescence images showing intensity of the ATF4 reporter, 4E-BP^intron^-DsRed in live wandering third instar larva from wildtype, *Hbs1* and *pelo* mutants. Scale bars represent 1μm.

**Fig. S2.**
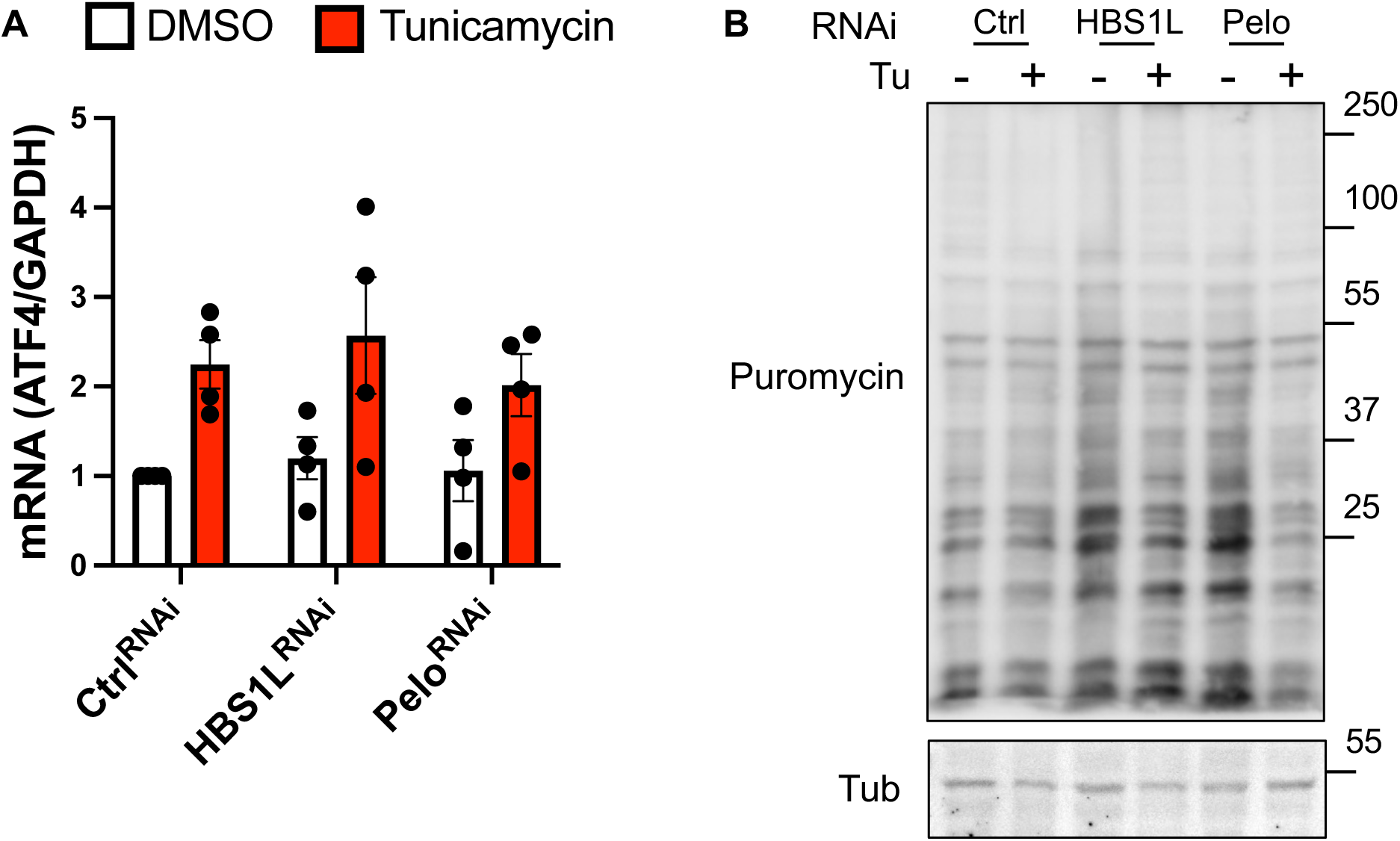
Depleting HBS1L or Pelo does not have an effect on ATF4 mRNA or global translation in HEK293T cells. **A.** q-RTPCR data measuring ATF4 mRNA normalized to house-keeping gene (GAPDH) in cells transfected with siRNA targeting HBS1L or Pelo and treated with 10μg/ml Tunicamycin to induce ER stress. Data represent mean of four independent experiments; error bars are standard error of mean. Statistical comparisons with Ctrl^RNAi^-DMSO were performed using the Wilcoxon signed rank test for and are shown as floating asterisks when significant. Comparisons with Ctrl^RNAi^- Tunicamycin were performed using the Friedman test with Dunn’s correction. Other comparisons were performed using the Wilcoxon matched pairs signed rank test. **B.** Representative western blot for puromycin-incorporation in cells where HBS1L or Pelo is depleted. Quantification presented in 2G.

**Fig. S3.**
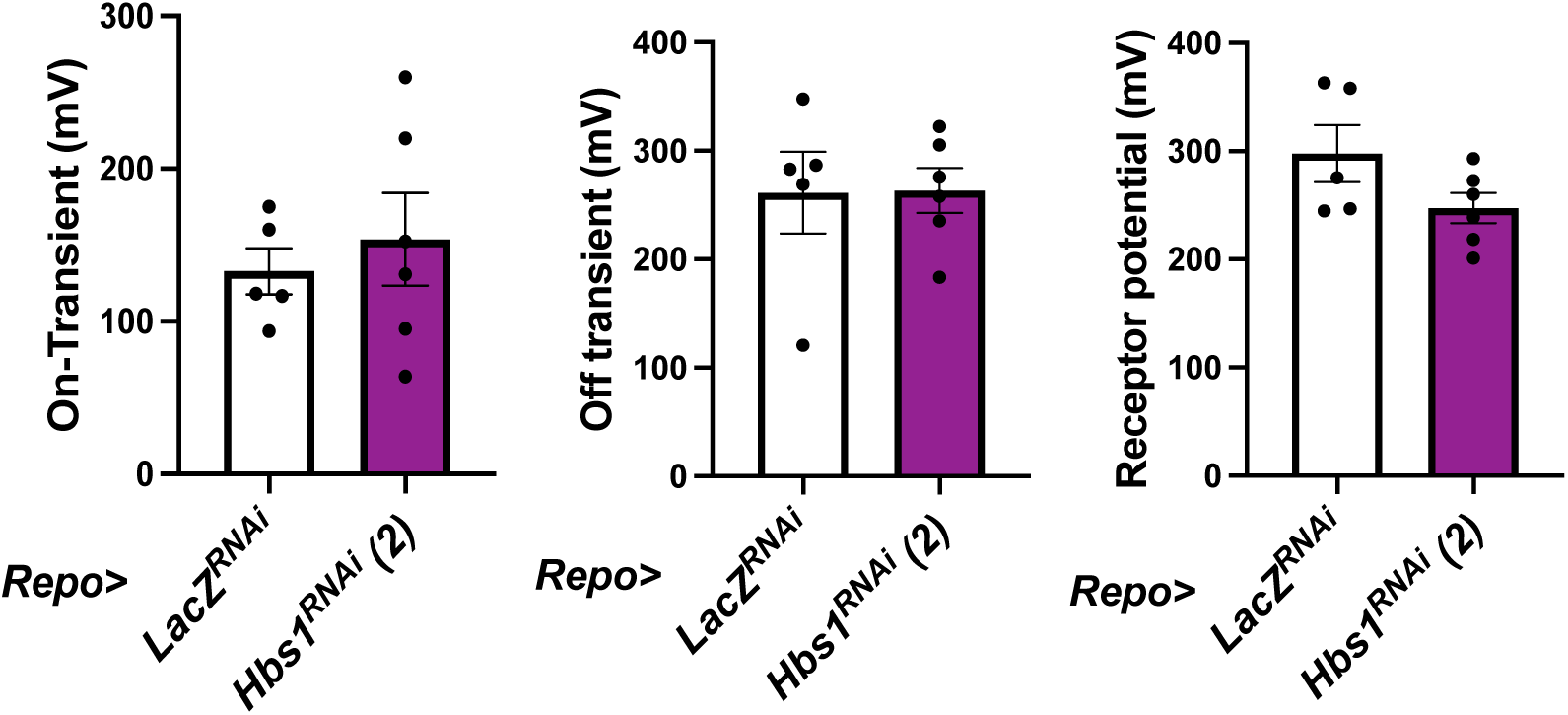
Depleting *Hbs1* in glia does not impact phototransduction. Quantification of on-transient, off-transient, and receptor potential from 3-5 day old animals where a glia driver (*Repo-GAL4*) is used to express a control *UAS-LacZ^RNAi^* or -*Hbs1^RNAi^*. Data are graphed as in 3E-G. Statistical comparisons were made using the Mann-Whitney test.

**Fig. S4.**
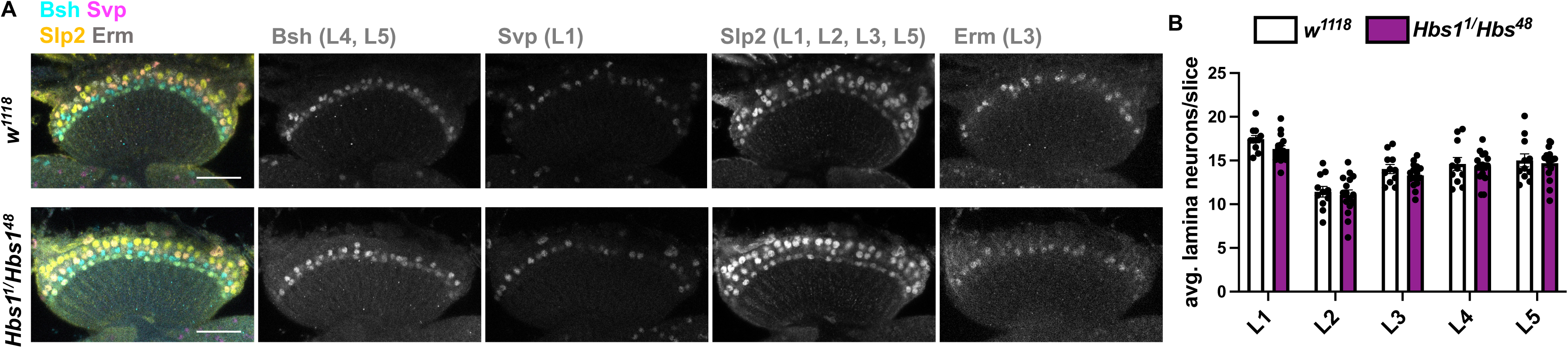
Loss of *Hbs1* does not alter lamina neuron number or morphology. **A.** Confocal images of 3-5 day old adult lamina from wildtype or *Hbs1* mutant animals stained markers for L1-L5 lamina neuron subtypes (Slp2 in cyan, Svp in magenta, Erm in yellow, and Bsh in gray). L1 neurons are Svp- and Slp2-positive, L2 neurons are only Slp2-positive, L3 are Erm-positive, L4 are only Bsh-positive, and L5 are Bsh- and Slp2-positive.

**Table S1.**
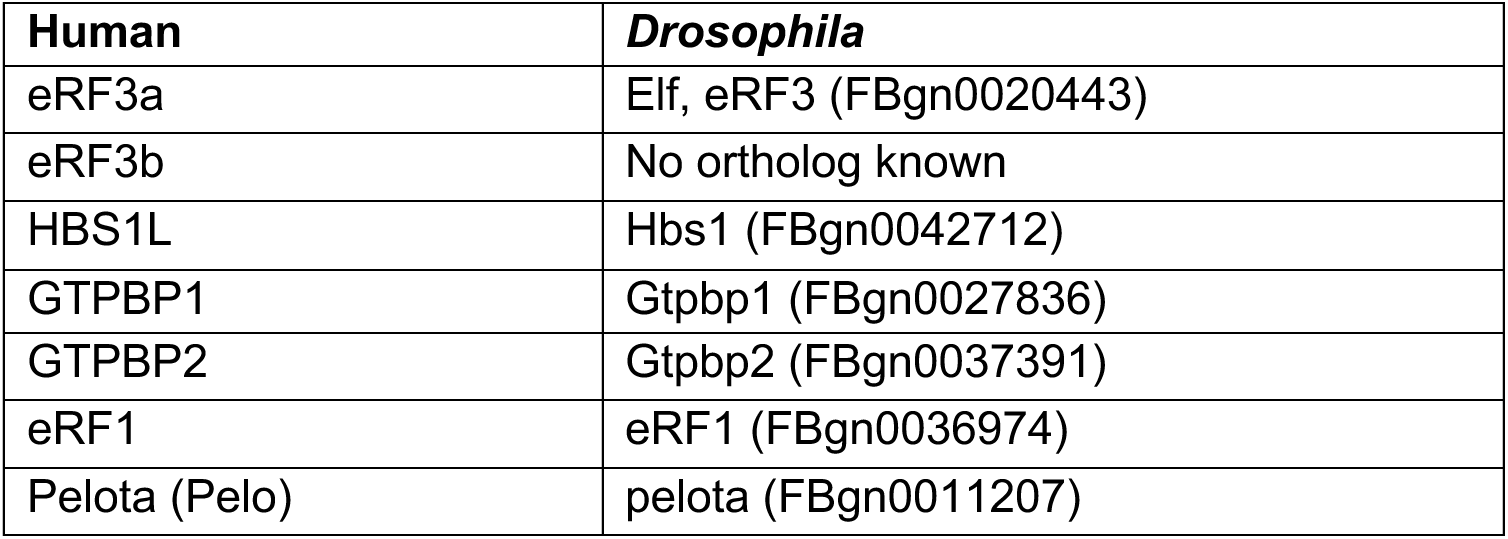
Human and corresponding *Drosophila* orthologs of genes encoding termination factors tested in the *Drosophila* RNAi screen. Where available, multiple RNAi lines per gene were used (see Table S2).

**Table S2.** *Drosophila* stocks used in this study.

| Transgene | Source | Figure |
| --- | --- | --- |
| *Dcg-GAL4* | BDSC_7011 | 1A-B |
| *4E-BP<sup>intron</sup>-DsRed* | (Kang et al. 2016) | 1A-F; S1 |
| *UAS-GFP* | BDSC_4776 | 1A |
| *UAS-LacZ<sup>RNAi</sup>* | (Kang et al. 2016) | 1A-B; 4A-E; 5A-B; S3 |
| *UAS-Hbs1<sup>RNAi</sup> (1)* | v104327 | 1A-B |
| *UAS-Hbs1<sup>RNAi</sup> (2)* | v33419 | 1A-B; 4A-E; S3 |
| *UAS-Elf<sup>RNAi</sup>* | v106240 |  |
| *UAS-pixie<sup>RNAi</sup> (1)* | v109630 |  |
| *UAS-pixie<sup>RNAi</sup> (2)* | v44325 |  |
| *UAS-Gtpbp1<sup>RNAi</sup> (1)* | v27490 |  |
| *UAS-Gtpbp1<sup>RNAi</sup> (2)* | v109410 |  |
| *UAS-Gtpbp2<sup>RNAi</sup>* | v107015 |  |
| *UAS-eRF1<sup>RNAi</sup>* | v45027 |  |
| *w<sup>1118</sup>* | BDSC_3605 | 1C-F; 3A, E-G; 6A-E; S1; S4 |
| *Hbs1<sup>1</sup>* | (Yang et al. 2015) | 1C-D; 3B-C, E-G; 6A-E; S1A; S4 |
| *Hbs1<sup>48</sup>* | (Yang et al. 2015) | 1C-D; 3B, E-G; 6A-E; S1A; S4 |
| *pelo<sup>1</sup>* | BDSC_11757 | 1E-F; 3C, E-G; S1B |
| *pelo<sup>PB60</sup>* | BDSC_68149 | 1E-F; S1B |
| *crc<sup>GFSTF</sup>* | BDSC_59608 | 3D-G |
| *Gcm-GAL4* | BDSC_45741 | 4A-E; 5A-H |
| *Repo-GAL4* | BDSC_7415 | S3 |
| *UAS-pelo<sup>RNAi</sup>* | v108606 | 5A-D |
| *UAS-ATF4<sup>RNAi</sup>* | v109014 | 5A-D |
| *UAS-LacZ* | BDSC_3956 | 5E-F |
| *UAS-ATF4(leaderless)* | (Vasudevan et al. 2020) | 5E-F |

## Notes

### Competing Interest Statement

The authors have declared no competing interest.

